# Phase-separated nuclear bodies of nucleoporin fusions, SET-NUP214 and NUP98-HOXA9, promote condensation of MLL1 and CRM1 to activate target genes

**DOI:** 10.1101/2022.05.24.493212

**Authors:** Masahiro Oka, Mayumi Otani, Yoichi Miyamoto, Jun Adachi, Takeshi Tomonaga, Munehiro Asally, Yasuyuki Ohkawa, Yoshihiro Yoneda

**Affiliations:** Laboratory of Nuclear Transport Dynamics, National Institutes of Biomedical Innovation, Health and Nutrition (NIBIOHN), Ibaraki, Osaka, Japan; Laboratory of Biomedical Innovation, Graduate School of Pharmaceutical Sciences, Osaka University, Suita, Osaka, Japan; Laboratory of Proteome Research and Laboratory of Proteomics for Drug Discovery, National Institutes of Biomedical Innovation, Health and Nutrition (NIBIOHN), Ibaraki, Osaka, Japans; School of Life Sciences, University of Warwick, United Kingdom; Department of Advanced Medical Initiatives, Faculty of Medicine, Kyushu University, Fukuoka, Japan; National Institutes of Biomedical Innovation, Health and Nutrition (NIBIOHN), Ibaraki, Osaka, Japan

**Keywords:** nucleoporin, NUP214, NUP98, MLL1, CRM1, leukemia, *HOX* cluster genes, nuclear body, phase separation, molecular condensation, gene expression

## Abstract

Nucleoporins NUP98 and NUP214 form chimeric fusion proteins that assemble into phase-separated nuclear bodies. However, the function and physiological significance of these nuclear bodies remain largely unknown. Previously, we reported that both NUP98-HOXA9 and SET-NUP214 are recruited to *HOX* cluster regions via chromatin-bound CRM1, a nuclear export receptor (Oka et al., 2019). Here, we show that these nuclear bodies promote the condensation of mixed lineage leukemia 1 (MLL1), a histone methyltransferase which is essential for the maintenance of *HOX* gene expression. Our analysis revealed that SET-NUP214 and CRM1 robustly associate with MLL1 to form nuclear bodies and are colocalized on chromatin. We also showed that MLL1 and CRM1 are recruited to the nuclear bodies of NUP98-HOXA9 and that the NUP98-HOXA9/CRM1/MLL1 complex accumulates on its target gene loci, including *HOX* clusters and *MEIS1*. These phenomena were not observed in phase-separation–deficient mutants or non-DNA-binding mutants of NUP98-HOXA9. Collectively, these results show that both phase separation and proper targeting of nucleoporin fusions to specific sites could enhance the activation of a wide range of target genes by promoting the condensation of MLL1 and CRM1.

## INTRODUCTION

NUP98 and NUP214 are nucleoporins, the components of the nuclear pore complex (NPC), and are often rearranged in leukemia (Lam and Aplan, 2001; Xu and Powers, 2009). The nucleoporin fusion genes, produced by chromosomal translocation, are associated with leukemogenesis: NUP98 fuses with various partner genes, including homeobox transcription factors (Gough et al., 2011; Michmerhuizen et al., 2020); NUP214 fuses with its partner genes, such as SET or DEK (Mendes and Fahrenkrog, 2019; Zhou and Yang, 2014).

Among NUP98 fusions, NUP98-HOXA9, a fusion between NUP98 and the homeobox transcription factor HOXA9 (Borrow et al., 1996; Nakamura et al., 1996), is among the best characterized fusion proteins to date. NUP98-HOXA9 causes aberrant gene expression that contributes to leukemogenesis (Kroon et al., 2001). Mechanistically, it is associated with epigenetic modifiers, such as CREB-binding protein (CBP)/p300, HDAC1, and MLL1 (Bai et al., 2006; Heikamp et al., 2021; Kasper et al., 1999; Mendes et al., 2020; Rio-Machin et al., 2017; Shima et al., 2017; Xu et al., 2016). In addition, NUP98 itself is known to be associated with trithorax (Trx), a Drosophila equivalent of MLL (Pascual-Garcia et al., 2014), or Wdr82– Set1A/COMPASS, another H3K4 methyltransferase (Franks et al., 2017). SET-NUP214, a fusion between NUP214 and the histone chaperone SET (von Lindern et al., 1992), also interacts with MLL1 or DOT1L (Cigdem et al., 2021; Van Vlierberghe et al., 2008). These studies suggest that the interaction between NUP fusion and these epigenetic modifiers likely plays an important role in triggering aberrant gene expression.

Intriguingly, NUP98 and NUP214 fusion proteins share three characteristics: (i) both contain dense FG repeats that are capable of forming nuclear bodies (Fornerod et al., 1995) through phase separation (Labokha et al., 2013). (Notably, recent studies demonstrated that the ability to form nuclear bodies of NUP98 fusion is important for aberrant gene activation and the transformation of hematopoietic cells (Ahn et al., 2021; Chandra et al., 2021; Terlecki-Zaniewicz et al., 2021)); (ii) their nuclear bodies colocalize with CRM1 (Oka et al., 2010; Oka et al., 2016; Port et al., 2016; Saito et al., 2016; Saito et al., 2004; Takeda et al., 2010), a nuclear export receptor (Fornerod et al., 1997; Fukuda et al., 1997; Ossareh-Nazari et al., 1997; Stade et al., 1997) originally identified in fission yeast (Adachi and Yanagida, 1989); (iii) both are associated with aberrant activation of *HOX* genes which encode the evolutionarily conserved transcription factors that function in various developmental processes (Krumlauf, 1994). It is known that *HOX* genes are dysregulated in various diseases and especially known to play a crucial role in leukemogenesis (Alharbi et al., 2013; Argiropoulos and Humphries, 2007; Gough et al., 2011; Grier et al., 2005; Hollink et al., 2011). Together, these studies suggest that the formation of nuclear bodies containing CRM1 is a key feature of leukemogenic NUP fusions. In support of this hypothesis, we recently found that CRM1 co-binds with NUP98- or NUP214-fusion proteins to chromatin at specific gene loci, including *HOX* clusters (Oka et al., 2019; Oka et al., 2016). In addition, CRM1 also facilitates the recruitment of SQSTM1-NUP214 (Gorello et al., 2010) to *HOX* genes (Lavau et al., 2020).

Of note, chromatin-bound CRM1 may also play a role in other types of leukemia. It has been shown that mutant NPM1 (NPM1c), the most frequent mutation in cytogenetically normal acute myeloid leukemia generating a novel nuclear export signal (NES) at its C-terminus (Falini et al., 2020; Falini et al., 2005; Schlenk et al., 2008), can activate *HOX* genes (Brunetti et al., 2018), and also co-binds with CRM1 to the *HOX* cluster region (Oka et al., 2019). Moreover, CALM-AF10, another NES-containing leukemogenic fusion protein, is also recruited to the *HOX* cluster via CRM1 (Conway et al., 2015).

Despite this evidence, the molecular mechanism by which the formation of nuclear bodies of NUP-fusion proteins linked to aberrant gene activation, especially its relationship with epigenetic modifiers, remains largely unknown.

In this study, we investigated the potential role of NUP-fusion nuclear bodies in the activation of its target genes. Our results suggested that the formation of NUP-fusion nuclear bodies, which is dependent on CRM1, is important for promoting the condensation of MLL1 on target sites to induce leukemogenic gene activation.

## RESULTS

### SET-NUP214 nuclear bodies colocalizes with HOX-B clusters

SET-NUP214 forms CRM1-containing nuclear bodies in the human T-ALL cell line LOUCY (Oka et al., 2019; Port et al., 2016). To evaluate the relevance of intranuclear localization of nuclear bodies of SET-NUP214, we performed immunoFISH. We focused on *HOX-B* clusters because our previous chromatin immunoprecipitation (ChIP)-seq data demonstrated that SET-NUP214 and CRM1 robustly accumulated at several specific loci, especially *HOX-B* clusters in LOUCY cells (Oka et al., 2019). Fluorescence microscopy images of *HOX-B* clusters and immunostaining of SET-NUP214 nuclear bodies showed that they overlap in the nucleus (Figure 1A, upper panel). To analyse this observation more quantitatively, we measured the shortest distance between *HOX-B* and SET-NUP214 for each nuclear bodies (n=478). This analysis revealed that 42.2 % of the nuclear bodies of SET-NUP214 were localized <0.2 μm from *HOX-B* clusters (Figure 1B, blue). As a negative control, we also performed immunoFISH for *RNF2* loci and the distance analysis (Figure 1A, lower, and Figure 1B, orange). Only 14.8 % of SET-NUP214 nuclear bodies were found within <0.2 μm from *RNF2* loci. These results, together with our previous ChIP-seq data, strongly suggest that the nuclear bodies of SET-NUP214/CRM1 preferentially colocalize with the *HOX-B* cluster regions.

**Figure 1.**
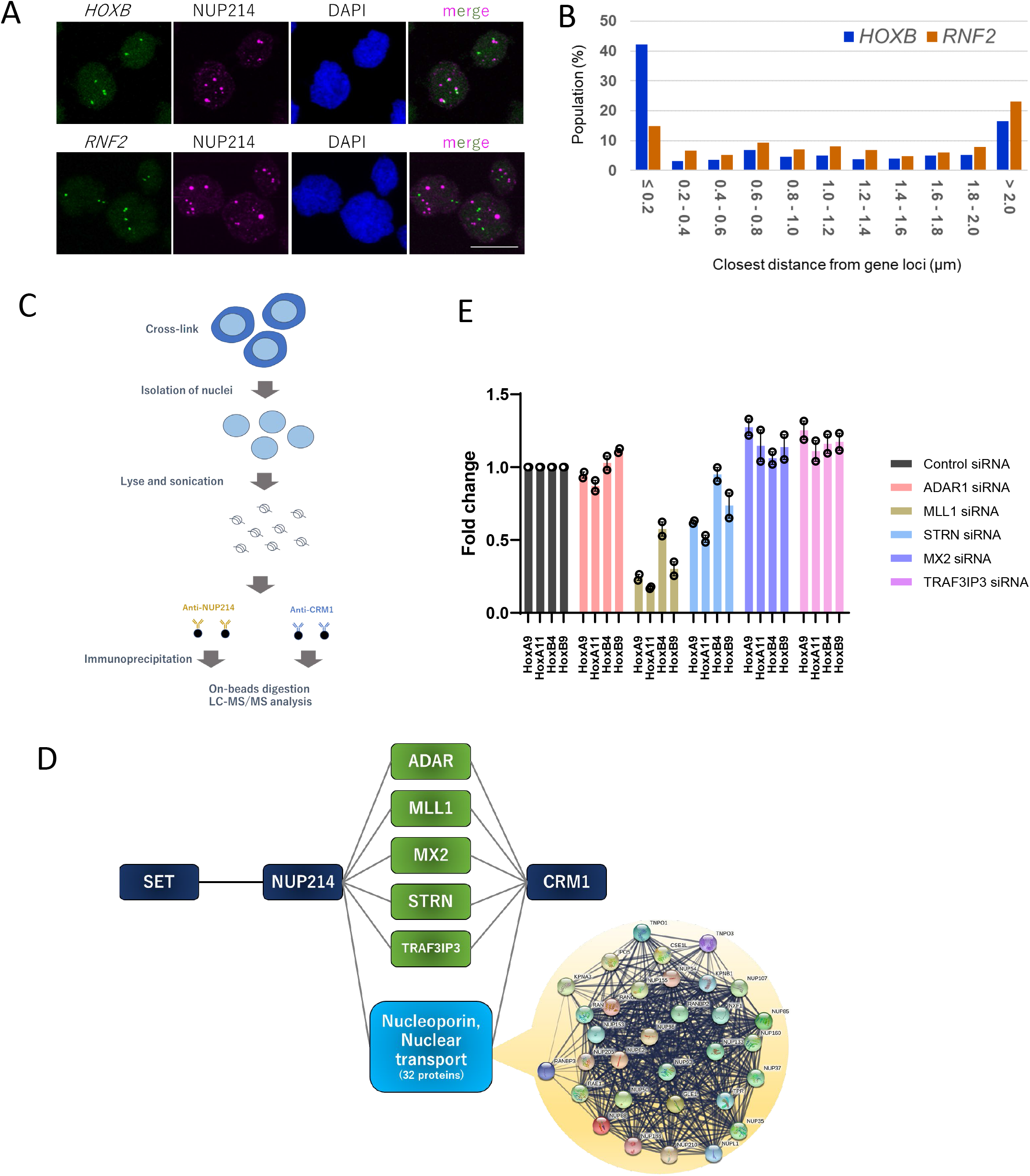
SET-NUP214 and CRM1 physically and functionally associate with MLL1. (A, B) Subcellular localization of SET-NUP214 and *HOX-B* or *RNF2* (control) gene loci in LOUCY cells. (A) ImmunoFISH was performed to detect SET-NUP214 nuclear bodies and respective gene loci. *HOX-B* cluster region is frequently found adjacent to SET-NUP214 nuclear bodies as compared with *RNF2*. Nuclei were stained with DAPI. Scale bars: 10 μm. (B) The minimum distance between each gene locus (as revealed by FISH) and the closest SET-NUP214 nuclear bodies was analyzed using ImageJ. (C) Schematic of RIME used in this study. (D) Summary of the RIME results. (E) Effect of knockdown of candidate components of SET-NUP214 nuclear bodies. Knockdown of ADAR1, KMT2A, STRN, MX2, and TARFIP3 was performed by nucleofection of siRNA (IDT) in LOUCY cells. The effect of knockdown on *HOX* gene expression (*HOXA9, HOXA11, HOXB4*, and *HOXB9*) was examined 4 days after nucleofection. *GAPDH* was used as the reference gene. The data are the average of two biological replicates each with two technical replicates.

### SET-NUP214/CRM1 physically and functionally associate with MLL1

Next, we attempted to identify the constituents of SET-NUP214/CRM1 nuclear bodies associated with chromatin. Taking advantage of the robust accumulation of SET-NUP214/CRM1 on *HOX* clusters in LOUCY cells, we performed the rapid immunoprecipitation mass spectrometry of endogenous proteins (RIME) (Figure 1C). This method uses formaldehyde to crosslink protein complexes and DNA, similar to ChIP-sequencing, and has been successfully used to identify chromatin-bound proteins associated with target molecules (Mohammed et al., 2013). Our RIME data demonstrated that NUP214 and CRM1 were strongly bound to each other, as their signals were primarily observed in both anti-NUP214 and anti-CRM1 immunoprecipitates (IPs) (Table I). As expected, SET was found in anti-NUP214 IPs, demonstrating that the SET-NUP214 fusion was successfully immunoprecipitated. To find out the factors that are involved in the function of SET-NUP214/CRM1 nuclear bodies, we focused on the proteins that are abundant both in anti-NUP214 IPs and anti-CRM1 IPs, but not in control IPs (strict criteria were used to select the candidate proteins whose peptide counts are either “0” or “1” in the control IP). We also found a majority of nucleoporins (22 nucleoporins), together with nuclear transport receptors (NTRs) or RAN and its regulators (Figure 1D, blue) in both IPs (Table I). We omitted these nucleoporins or proteins involved in nuclear transport from our analysis since they are most likely the association with endogenous NUP214 or CRM1 on NPCs. Interestingly, the analysis revealed several proteins whose relationship with NPC or nuclear transport is not known to date, but are strongly associated with both NUP214 and CRM1: namely, ADAR, MLL1, striatin, and TRAF3IP(T3JAM) (Figure 1D). We also found MX2, an antiviral protein that interacts with several nucleoporins (Dicks et al., 2018). We further analysed these five proteins.

**Table I.**
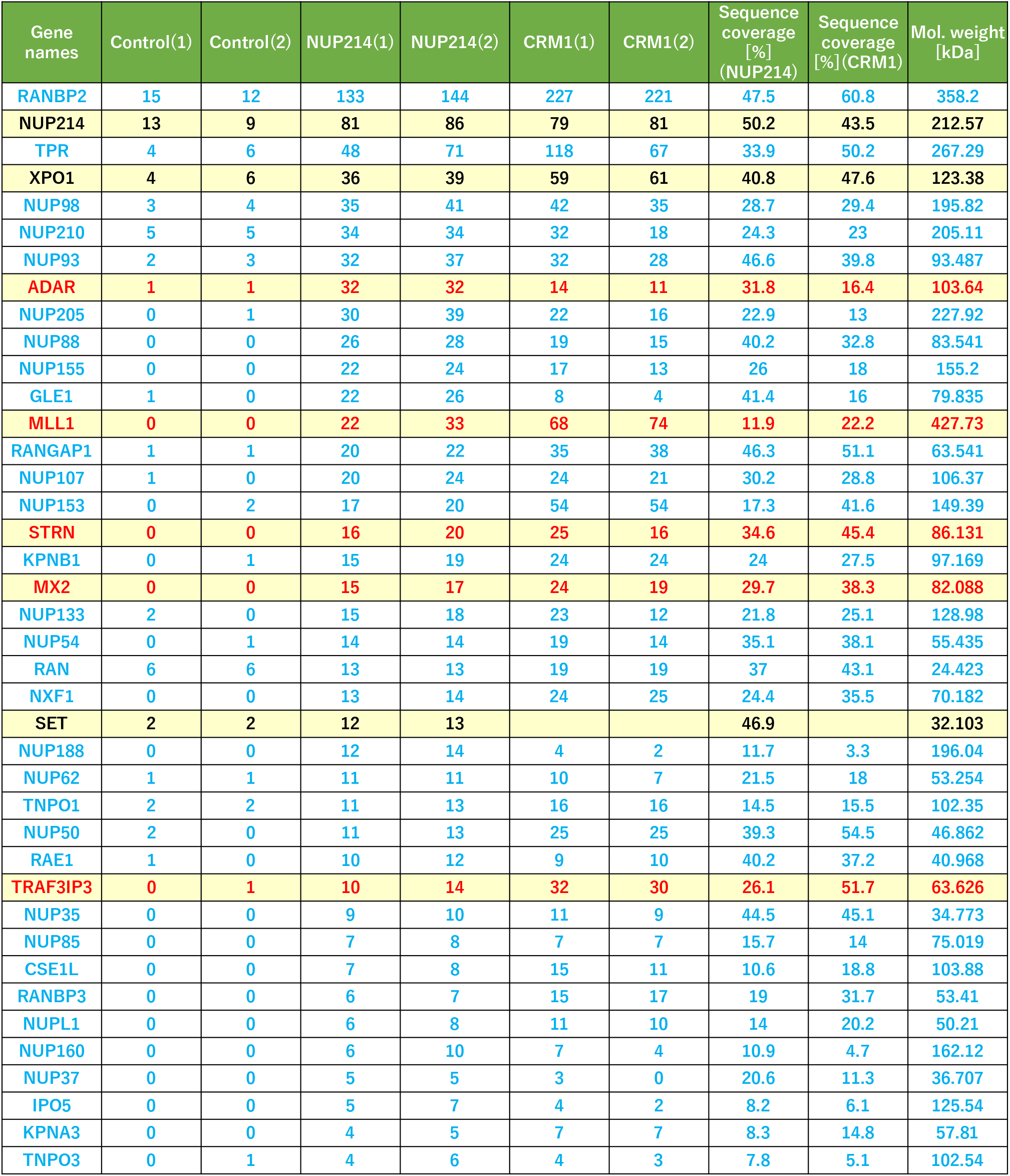
SET-NUP214 or CRM1 associated proteins in LOUCY cells were identified using RIME. For each target protein, RIME was performed in duplicate using specific antibodies or control IgG, and the number of peptides identified and sequence coverage are shown. Nucleoporins and nuclear transport-related proteins are indicated by blue letters. Several proteins (red letters) are not involved in the nuclear transport activity.

| Gene names | Control(1) | Control(2) | NUP214(1) | NUP214(2) | CRM1(1) | CRM1(2) | Sequence coverage [%] (NUP214) | Sequence coverage [%](CRM1) | Mol. weight [kDa] |
| --- | --- | --- | --- | --- | --- | --- | --- | --- | --- |
| RANBP2 | 15 | 12 | 133 | 144 | 227 | 221 | 47.5 | 60.8 | 358.2 |
| NUP214 | 13 | 9 | 81 | 86 | 79 | 81 | 50.2 | 43.5 | 212.57 |
| TPR | 4 | 6 | 48 | 71 | 118 | 67 | 33.9 | 50.2 | 267.29 |
| XPO1 | 4 | 6 | 36 | 39 | 59 | 61 | 40.8 | 47.6 | 123.38 |
| NUP98 | 3 | 4 | 35 | 41 | 42 | 35 | 28.7 | 29.4 | 195.82 |
| NUP210 | 5 | 5 | 34 | 34 | 32 | 18 | 24.3 | 23 | 205.11 |
| NUP93 | 2 | 3 | 32 | 37 | 32 | 28 | 46.6 | 39.8 | 93.487 |
| ADAR | 1 | 1 | 32 | 32 | 14 | 11 | 31.8 | 16.4 | 103.64 |
| NUP205 | 0 | 1 | 30 | 39 | 22 | 16 | 22.9 | 13 | 227.92 |
| NUP88 | 0 | 0 | 26 | 28 | 19 | 15 | 40.2 | 32.8 | 83.541 |
| NUP155 | 0 | 0 | 22 | 24 | 17 | 13 | 26 | 18 | 155.2 |
| GLE1 | 1 | 0 | 22 | 26 | 8 | 4 | 41.4 | 16 | 79.835 |
| MLL1 | 0 | 0 | 22 | 33 | 68 | 74 | 11.9 | 22.2 | 427.73 |
| RANGAP1 | 1 | 1 | 20 | 22 | 35 | 38 | 46.3 | 51.1 | 63.541 |
| NUP107 | 1 | 0 | 20 | 24 | 24 | 21 | 30.2 | 28.8 | 106.37 |
| NUP153 | 0 | 2 | 17 | 20 | 54 | 54 | 17.3 | 41.6 | 149.39 |
| STRN | 0 | 0 | 16 | 20 | 25 | 16 | 34.6 | 45.4 | 86.131 |
| KPNB1 | 0 | 1 | 15 | 19 | 24 | 24 | 24 | 27.5 | 97.169 |
| MX2 | 0 | 0 | 15 | 17 | 24 | 19 | 29.7 | 38.3 | 82.088 |
| NUP133 | 2 | 0 | 15 | 18 | 23 | 12 | 21.8 | 25.1 | 128.98 |
| NUP54 | 0 | 1 | 14 | 14 | 19 | 14 | 35.1 | 38.1 | 55.435 |
| RAN | 6 | 6 | 13 | 13 | 19 | 19 | 37 | 43.1 | 24.423 |
| NXF1 | 0 | 0 | 13 | 14 | 24 | 25 | 24.4 | 35.5 | 70.182 |
| SET | 2 | 2 | 12 | 13 |  |  | 46.9 |  | 32.103 |
| NUP188 | 0 | 0 | 12 | 14 | 4 | 2 | 11.7 | 3.3 | 196.04 |
| NUP62 | 1 | 1 | 11 | 11 | 10 | 7 | 21.5 | 18 | 53.254 |
| TNPO1 | 2 | 2 | 11 | 13 | 16 | 16 | 14.5 | 15.5 | 102.35 |
| NUP50 | 2 | 0 | 11 | 13 | 25 | 25 | 39.3 | 54.5 | 46.862 |
| RAE1 | 1 | 0 | 10 | 12 | 9 | 10 | 40.2 | 37.2 | 40.968 |
| TRAF3IP3 | 0 | 1 | 10 | 14 | 32 | 30 | 26.1 | 51.7 | 63.626 |
| NUP35 | 0 | 0 | 9 | 10 | 11 | 9 | 44.5 | 45.1 | 34.773 |
| NUP85 | 0 | 0 | 7 | 8 | 7 | 7 | 15.7 | 14 | 75.019 |
| CSE1L | 0 | 0 | 7 | 8 | 15 | 11 | 10.6 | 18.8 | 103.88 |
| RANBP3 | 0 | 0 | 6 | 7 | 15 | 17 | 19 | 31.7 | 53.41 |
| NUPL1 | 0 | 0 | 6 | 8 | 11 | 10 | 14 | 20.2 | 50.21 |
| NUP160 | 0 | 0 | 6 | 10 | 7 | 4 | 10.9 | 4.7 | 162.12 |
| NUP37 | 0 | 0 | 5 | 5 | 3 | 0 | 20.6 | 11.3 | 36.707 |
| IPO5 | 0 | 0 | 5 | 7 | 4 | 2 | 8.2 | 6.1 | 125.54 |
| KPNA3 | 0 | 0 | 4 | 5 | 7 | 7 | 8.3 | 14.8 | 57.81 |
| TNPO3 | 0 | 1 | 4 | 6 | 4 | 3 | 7.8 | 5.1 | 102.54 |

Knockdowns of the five candidate genes were performed in LOUCY cells to monitor the effect on the expression of *HOX* genes, which is dependent on SET-NUP214. The fold change in *HOX* genes’ expressions were measured in LOUCY cells by qPCR (Figure 1E). We found that the knockdown of MLL1 substantially decreased the expression of *HOX* genes (Figure 1E, Figure 1—figure supplement 1). We also found that knockdown of striatin, a scaffold protein involved in the regulation of signaling pathways (Hwang and Pallas, 2014), also decreased the expression of *HOX* genes. The knockdown of other candidate genes did not affect *HOX* gene expression. Since the effect of MLL1 knockdown was more evident than that of other knockdowns, we focused our analysis on MLL1.

To characterize MLL1, we costained the LOUCY cells with anti-MLL1 and anti-CRM1 antibodies (Figure 2, upper two columns). MLL1 frequently colocalized with SET-NUP214/CRM1 nuclear bodies in LOUCY cells (Figure 2), which is consistent with recent report (Cigdem et al., 2021). To examine if the partner proteins of MLL1 also colocalize with the nuclear bodies, we immunostained Menin which has been shown to interact with MLL1 physically and functionally (Matkar et al., 2013; Yokoyama et al., 2005). The immunstaining showed that Menin also colocalize with SET-NUP214/CRM1 nuclear bodies (Figure 2, third column). In contrast, BRD4, a super-enhancer (SE)-enriched transcriptional coactivator that often forms nuclear puncta (Sabari et al., 2018), only partially colocalized with SET-NUP214 nuclear bodies.

**Figure 2.**
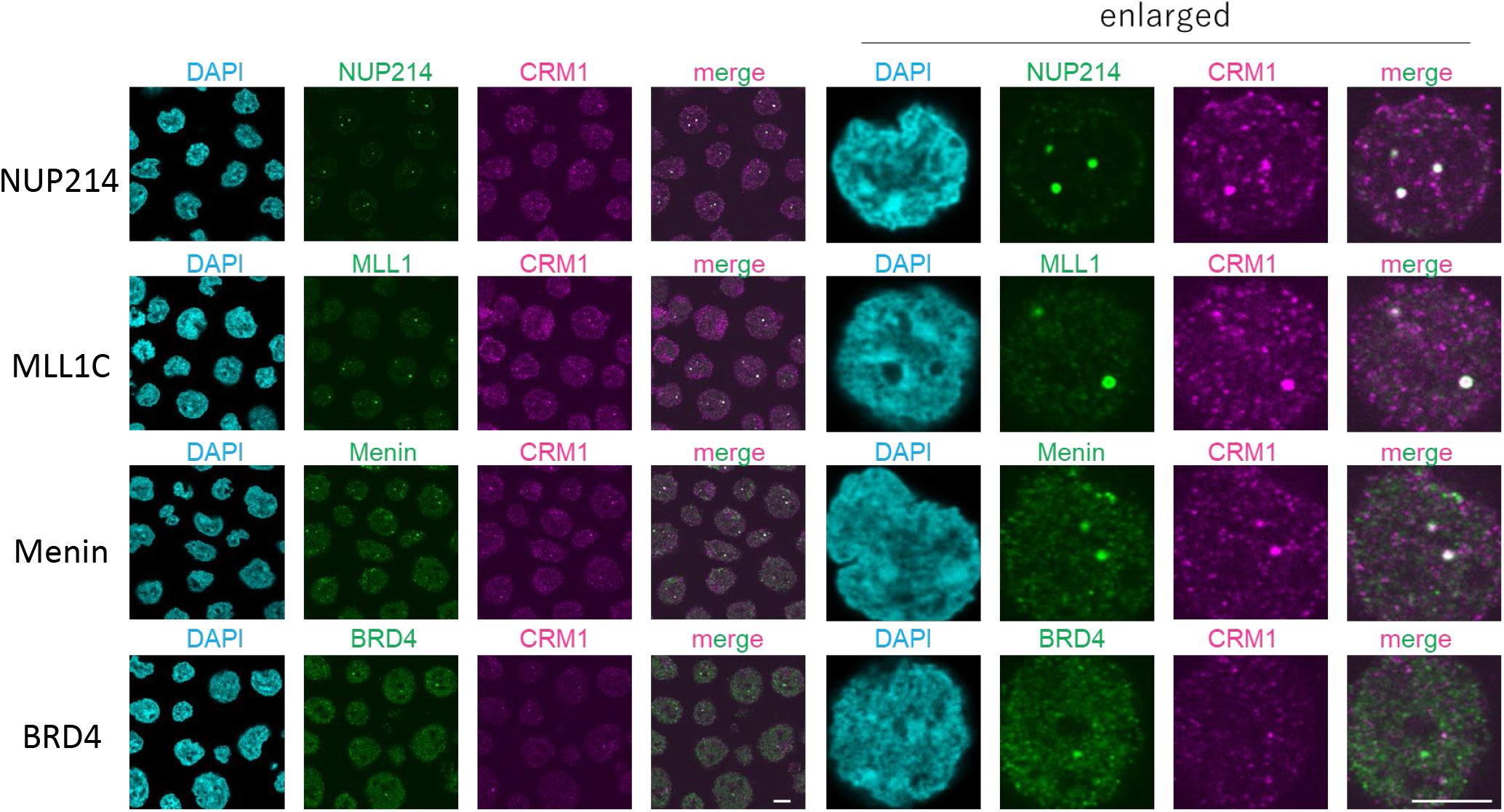
SET-NUP214 nuclear bodies colocalized with CRM1, MLL1 and Menin. LOUCY cells were immunostained with anti-NUP214, MLL1C, Menin, BRD4, and CRM1 antibodies. The merged images of CRM1 are shown. Nuclei were stained with DAPI. Scale bar: 5 μm.

### SET-NUP214/CRM1/MLL1 show genome-wide colocalization and are essential constituents of the nuclear bodies

To further characterise the relevance of the association between MLL1, SET-NUP214, and CRM1 in genome-wide binding, we performed ChIP-seq analysis (Figure 3A). Strikingly, the binding sites of these proteins showed frequent co-occupancy on a genome-wide scale, including *HOX-A* and *HOX-B* cluster regions (Figure 3A and 3B). Next, we examined the role of MLL1 in SET-NUP214/CRM1 accumulation in *HOX* clusters. ChIP-qPCR analysis revealed that MLL1 knockdown caused a significant decrease in SET-NUP214 signals in *HOX* clusters (Figure 3C). Furthermore, CRM1 signals in the *HOX* clusters were significantly diminished. These results suggested that binding of both SET-NUP214 and CRM1 to *HOX* cluster regions is maintained by MLL1 and implicate that SET-NUP214, CRM1, and MLL1 form functional nuclear bodies associated with *HOX* cluster regions to robustly activate *HOX* genes.

**Figure 3.**
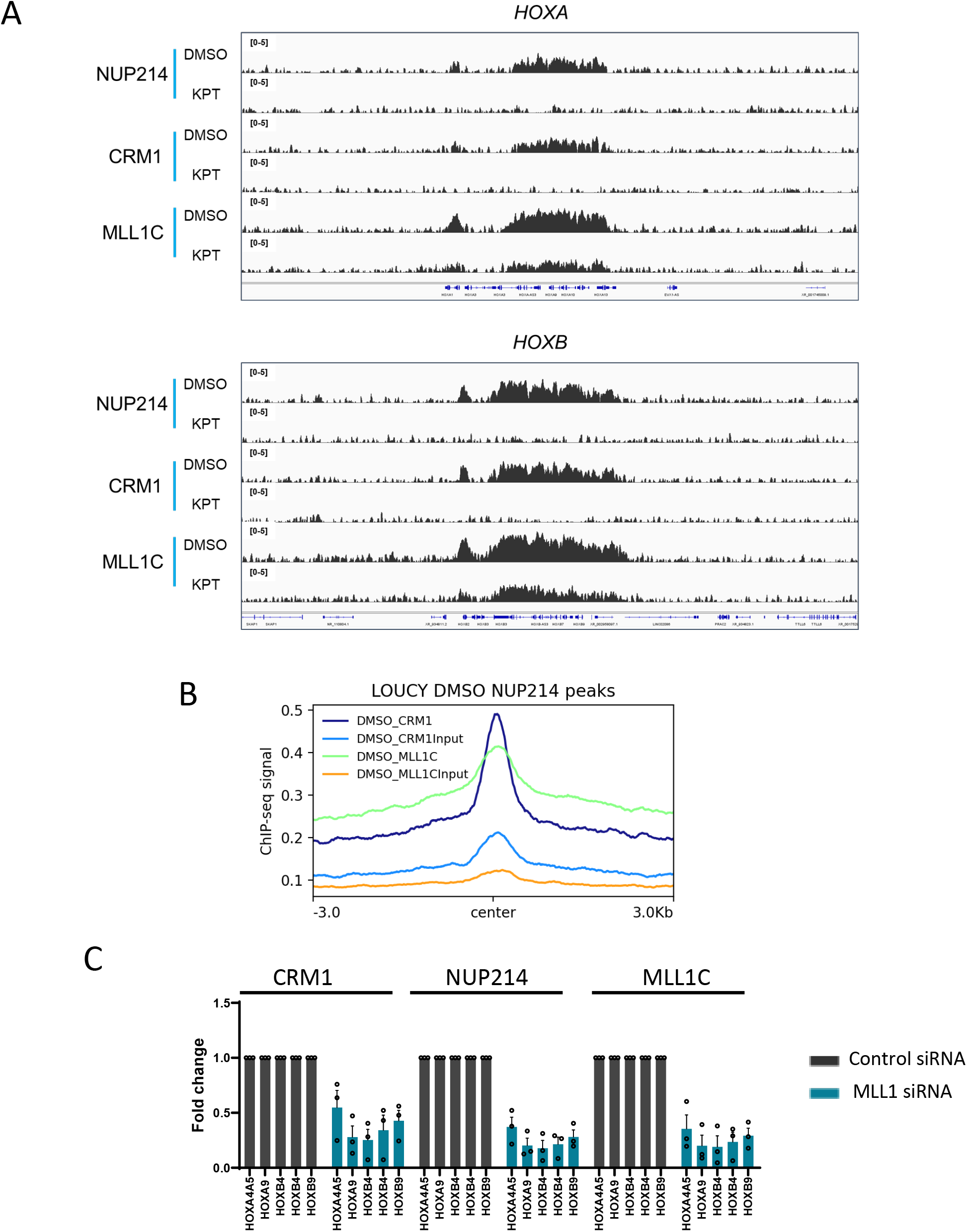
MLL1 is an essential component of SET-NUP214 nuclear bodies to recruit and activate target genes. (A) The binding profiles of NUP214, CRM1, and MLL1C in LOUCY cells treated either in the presence of DMSO (vehicle control) or KPT-330 (1,000 nM) for 24 h (ChIP-seq data for NUP214 and CRM1 were from previous study: GSE127983.) (B) Aggregation plots of CRM1 and MLL1C binding sites in LOUCY. CRM1 binding signals and MLL1C binding signals are mapped against NUP214 binding sites. (C) ChIP-qPCR analysis of CRM1, NUP214, and MLL1 at *HOX* gene loci in LOUCY cells treated either with control siRNA or MLL1 siRNA for 4 days. The primer set used was as follows: HOXA4A5 (intergenic region between *HOXA4* and *HOXA5*); HOXA9 (promoter); HOXB4 (promoter); HOXB4 (enhancer); HOXB9 (promoter). Data are presented as mean values ± SEM of three independent experiments (n = 3).

To investigate the role of CRM1 in this process, we used the CRM1 inhibitor KPT-330, which covalently binds to CRM1 to inhibit its export activity and induces its degradation (Etchin et al., 2013; Lapalombella et al., 2012; Mendonca et al., 2014; Tai et al., 2014; Turner et al., 2013; Zhang et al., 2013) and disassembly of SET-NUP214 nuclear bodies (Oka et al., 2019). First, immunostaining revealed that SET-NUP214 nuclear bodies colocalizing with MLL1 diminished when incubated with 100 nM of KPT-330 for 24 h, and those were almost completely disrupted at 1 μM (Figure 4A). Next, ChIP-seq and ChIP-qPCR were performed to monitor the effect of KPT-330 treatment on the chromatin binding of MLL1 (Figure 3A, 4B). The results showed that KPT-330 treatment caused a significant decrease in the signals of MLL1 in *HOX* cluster regions, which is similar to the results observed in both CRM1 and SET-NUP214. We also monitored the effect of KPT-330 on the MLL protein levels. As shown in Figure 4C, our results showed that MLL1 protein levels were not significantly affected by KPT-330. In addition, LMB, a CRM1 inhibitor that also disassembles nuclear bodies (Port et al., 2016; Saito et al., 2016) and destabilizes the SET-NUP214 protein (Oka et al., 2019), without inducing the degradation of CRM1, did not affect the levels of MLL1. Collectively, these results suggest that CRM1 is critical for the assembly of SET-NUP214 nuclear bodies that can recruit MLL1.

**Figure 4.**
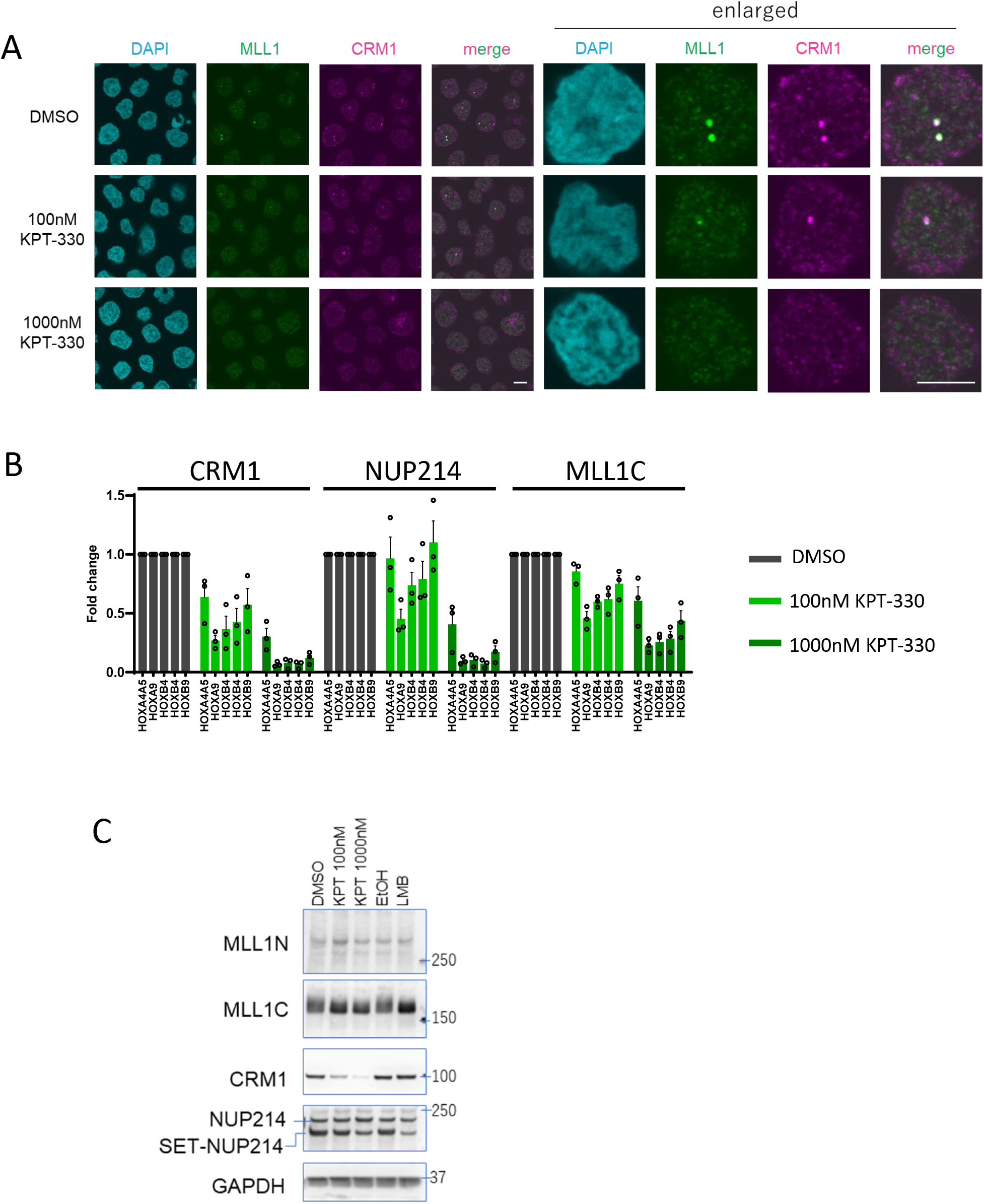
The effect of CRM1 inhibitor on the function of SET-NUP214 nuclear bodies. (A) LOUCY cells treated with DMSO, 100 or 1,000 nM of KPT-330 for 24 h were co-immunostained with anti-CRM1 and MLL1C antibodies. DAPI staining was used to visualize nuclei. Scale bar: 5 μm. (B) ChIP-qPCR analysis of CRM1, NUP214, and MLL1C at *HOX* gene loci in LOUCY cells cultured either in the presence of DMSO (vehicle control) or KPT-330 (1,000 nM) for 24 h. Data are presented as mean values ± SEM of three independent experiments (n = 3). (C) Protein expression levels of MLL1N, MLL1C, CRM1, SET-NUP214, and GAPDH in LOUCY cells treated with DMSO (vehicle control for KPT-330) or KPT-330 (100 or 1,000 nM), EtOH (vehicle control for LMB) or LMB (10 nM) for 24 h. The cell lysates were prepared in RIPA buffer and analyzed by immunoblotting.

### Phase separation and DNA binding of NUP98-HOXA9 are important for its targeting and downstream gene activation

Previous reports suggested unexpected similarities between the nuclear bodies formed by SET-NUP214 and NUP98-HOXA9; namely, they both accumulate on *HOX* cluster regions (Oka et al., 2019; Oka et al., 2016; Van Vlierberghe et al., 2008; Xu et al., 2016) and contain CRM1 (Oka et al., 2010; Oka et al., 2016; Saito et al., 2004; Takeda et al., 2010). NUP98-HOXA9 is also known to be physically and functionally associated with MLL1 in hematopoietic progenitor cells (Shima et al., 2017; Xu et al., 2016). However, the relevance of the nuclear bodies of NUP98-HOXA9 to MLL1 remains unknown. These results prompted us to investigate the relationship between MLL1 and the nuclear bodies formed by NUP98-HOXA9.

To investigate the function of NUP98-HOXA9, we isolated mouse ES cells (mESCs) stably expressing FLAG-tagged NUP98-HOXA9 since we demonstrated that the *Hox* genes are selectively activated in these cells (Oka et al., 2016). We first confirmed that our newly established cell lines robustly activated *Hox* genes (Figure 5A). This analysis revealed that other targets of NUP98-HOXA9 in transformed leukemia cells, *Meis1* and *Pbx3* (Takeda et al., 2006; Xu et al., 2016), were also activated in these cells. This finding suggests that the mechanism of NUP98-HOXA9-mediated gene activation is highly conserved between mESCs and transformed leukemia cells.

**Figure 5.**
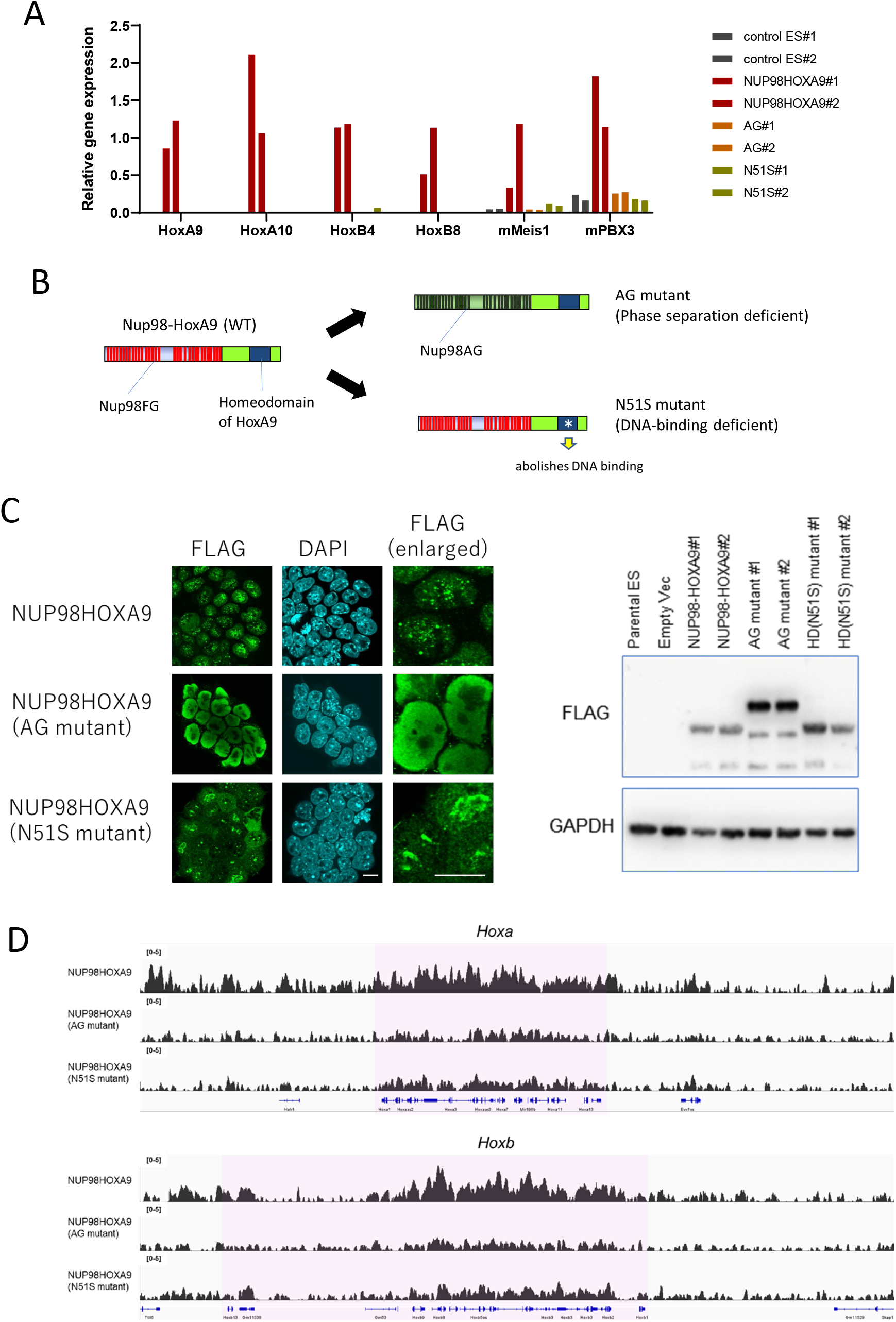
FG repeat and homeodomain of NUP98-HOXA9 is important for its accumulation on HOX clusters and gene activation. (A) qPCR analysis of NUP98-HOXA9-target genes (*Hoxa9, Hoxa10, Hoxb4, Hoxb8, Meis1, Pbx3*) in indicated NUP98-HOXA9 expressing stable cell lines. Two independent cell lines for each (control, wild-type NUP98-HOXA9, AG mutant, and N51S homeodomain mutant) were analyzed. *Gapdh* was used as a reference gene. (B) Schematic representation of NUP98-HOXA9, FG mutant, or homeodomain (N51S) mutant. (C) (left) Confocal imaging of FLAG-NUP98-HOXA9, AG mutant, or N51S mutant. DAPI staining was used to visualize the nuclei. (right) Protein expression levels of FLAG-NUP98-HOXA9 and its mutants. GAPDH was used as a loading control. (D) Binding profiles of FLAG-NUP98-HOXA9, AG mutant, or N51S mutant in mESC stable cell lines.

To determine the importance of chromatin-associated nuclear bodies formed by NUP98-HOXA9, we established cell lines carry mutated versions of NUP98-HOXA9. AG mutant (mutated from FG to AG) is deficient in phase separation and N51S mutant (possessing a point mutation in the homeodomain [N51S](Shanmugam et al., 1999)) is deficient to bind DNA (Figure 5B). Introducing the AG mutation completely abolished the formation of nuclear bodies, while N51S did not abolish the formation of nuclear bodies. However, these were fewer and larger than that formed by wild-type NUP98-HOXA9 (Figure 5C), which is consistent with previous studies (Ahn et al., 2021; Fahrenkrog et al., 2016).

These mutants enabled us to examine the importance of the formation of nuclear bodies or their DNA-binding on accumulation in the *Hox* cluster region and gene activation. Specifically, we compared ChIP data between NUP98-HOXA9 and its mutants. Our ChIP-seq and qPCR results showed that the formation of nuclear bodies by FG repeats was important for both the accumulation on *Hox* cluster genes and their robust activation (Figure 5A, D). In AG mutant, the signals on *Hox* clusters were significantly diminished, and *Hox* gene activation was completely lost. Thus, phase separation of NUP98-HOXA9 significantly contributes to its targeting and concomitant gene activation. These results agree with a recent report using similar mutant construct (FG to SG) expressed in hematopoietic stem and progenitor cells (Ahn et al., 2021). Our results also showed that DNA-binding of nuclear bodies is important for the targeting of NUP98-HOXA9, since the N51S mutation caused a drastic decrease in the ChIP-signal on *Hox* clusters and terminated the activation of *Hox* genes.

Intriguingly, these phenomena were observed for target genes other than *Hox* clusters, such as *Meis1* or *Pbx3*, in mESCs. Collectively, these results showed that the phase separation and DNA-binding of NUP98-HOXA9 nuclear bodies, two distinct properties of the fusion that rely on FG repeats and homeodomain, respectively, are important for targeting nuclear bodies onto *Hox* clusters and other targets to activate downstream genes.

### NUP98-HOXA9 nuclear bodies induce the condensation of CRM1 and MLL1 on its target loci

Using these cell lines, we examined whether the nuclear bodies formed by NUP98-HOXA9 are also related to MLL1. Strikingly, immunofluorescence analysis revealed that most of the nuclear bodies formed by FLAG-NUP98-HOXA9 colocalized with MLL1 (Figure 6A). We also noticed that these unusual staining patterns of MLL (obvious speckle formation) were only observed in NUP98-HOXA9-expressing ES cells but not in parental ES cells, AG mutants, or N51S homeodomain mutant expressing ES cells. Note that even if the robust NUP98-HOXA9 nuclear bodies are formed in N51S mutant-expressing cells, MLL1 is not recruited in these nuclear bodies. Therefore, our results suggested that both phase separation (via FG repeats) and proper targeting to specific genome loci (via functional homeodomain) are necessary for the formation of NUP98 nuclear bodies that could induce the sequestration of MLL1. These results showed that nuclear bodies formed by NUP98-HOXA9 induce molecular condensation of MLL1, which may increase the local concentration of MLL on *Hox* clusters or other target loci.

**Figure 6.**
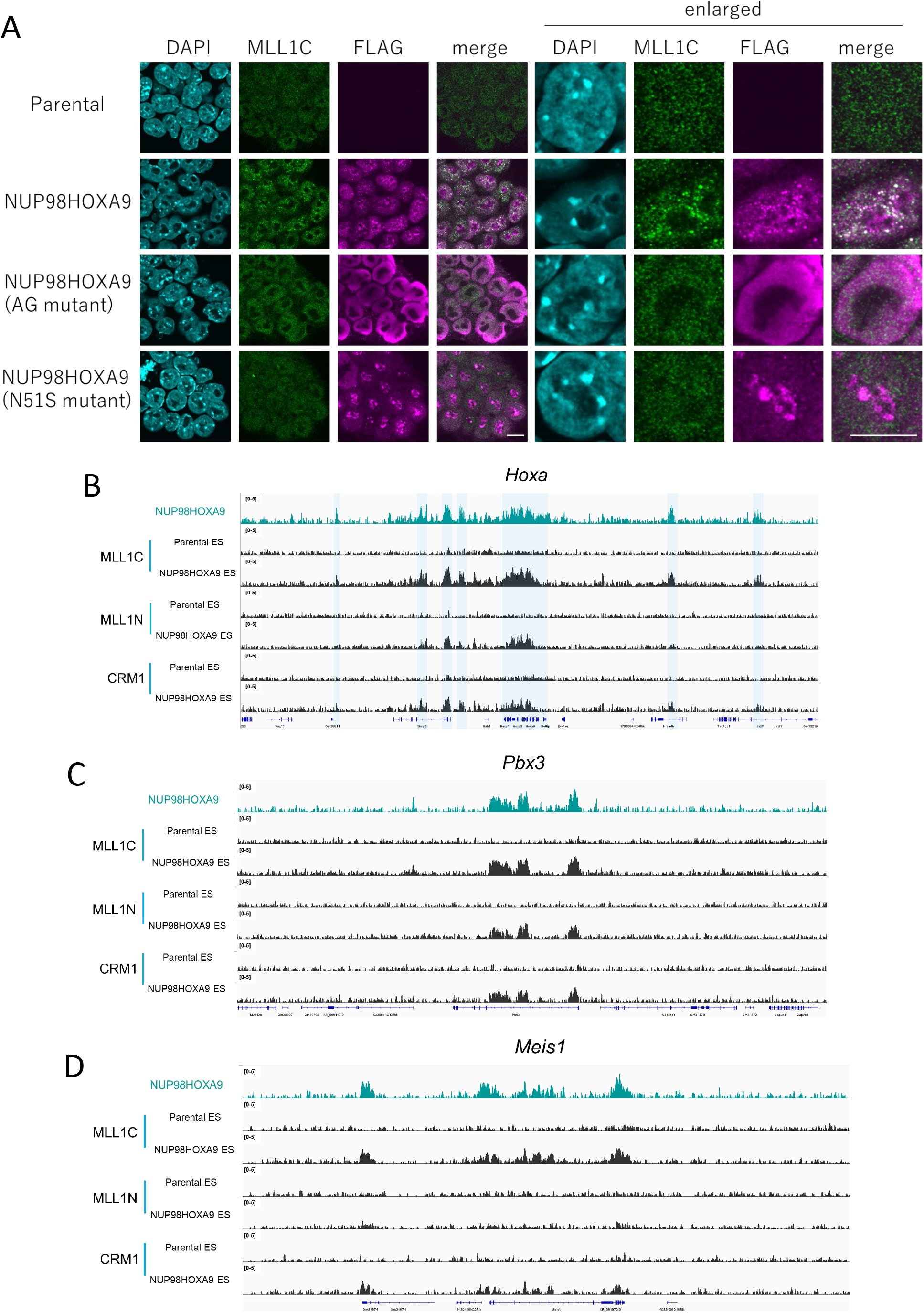
NUP98-HOXA9 nuclear bodies stimulate the condensation of MLL1 and its accumulation on target sites. (A) Subcellular localization of MLL1C in stable mESCs. Parental mESCs or its stable cell lines expressing FLAG-NUP98-HOXA9, AG mutant, homeodomain mutant, were co-immunostained with anti-MLL1C and FLAG. DAPI staining for nuclei. (B–D) The binding profiles of MLL1C, MLL1N, CRM1 in parental mESC or stable cell lines expressing FLAG-NUP98-HOXA9 as revealed with ChIP-seq (B:*Hoxa*, C:*Pbx3*, D:*Meis1*).

To further investigate the role of NUP98-HOXA9 in the accumulation of MLL1 at the target sites, we performed ChIP-seq analysis (Figure 6B–D, Figure 6—figure supplement 1). We found that MLL1 (both MLL1N and MLL1C) weakly bound to the *Hox* cluster and other target regions in parental ES cells. However, MLL1 robustly accumulated on the target sites in the cell line expressing FLAG-NUP98-HOXA9. As for CRM1, we previously demonstrated that ChIP-seq peaks of CRM1 were significantly enhanced in FLAG-NUP98-HOXA9 expressing cells as compared with parental ES cells, using polyclonal anti-CRM1 antibody (Oka et al., 2016). Here, we noticed that ChIP-seq using monoclonal anti-CRM1 antibody (CST) showed marked differences in CRM1 ChIP signals between parental ES cells and FLAG-NUP98-HOXA9 expressing cells (Figure 6B–D).

We further performed ChIP-qPCR and found that these effects were only observed in the *Hox* cluster region in wild-type NUP98-HOXA9-expressing cells but not in AG or N51S mutants (Figure 7A). In addition, correlation plot showed that there were strong positive correlations between the binding sites of CRM1 and MLL1 in parental ES cells, and those were significantly enhanced by the expression of FLAG-NUP98-HOXA9 (Figure 7B).

**Figure 7.**
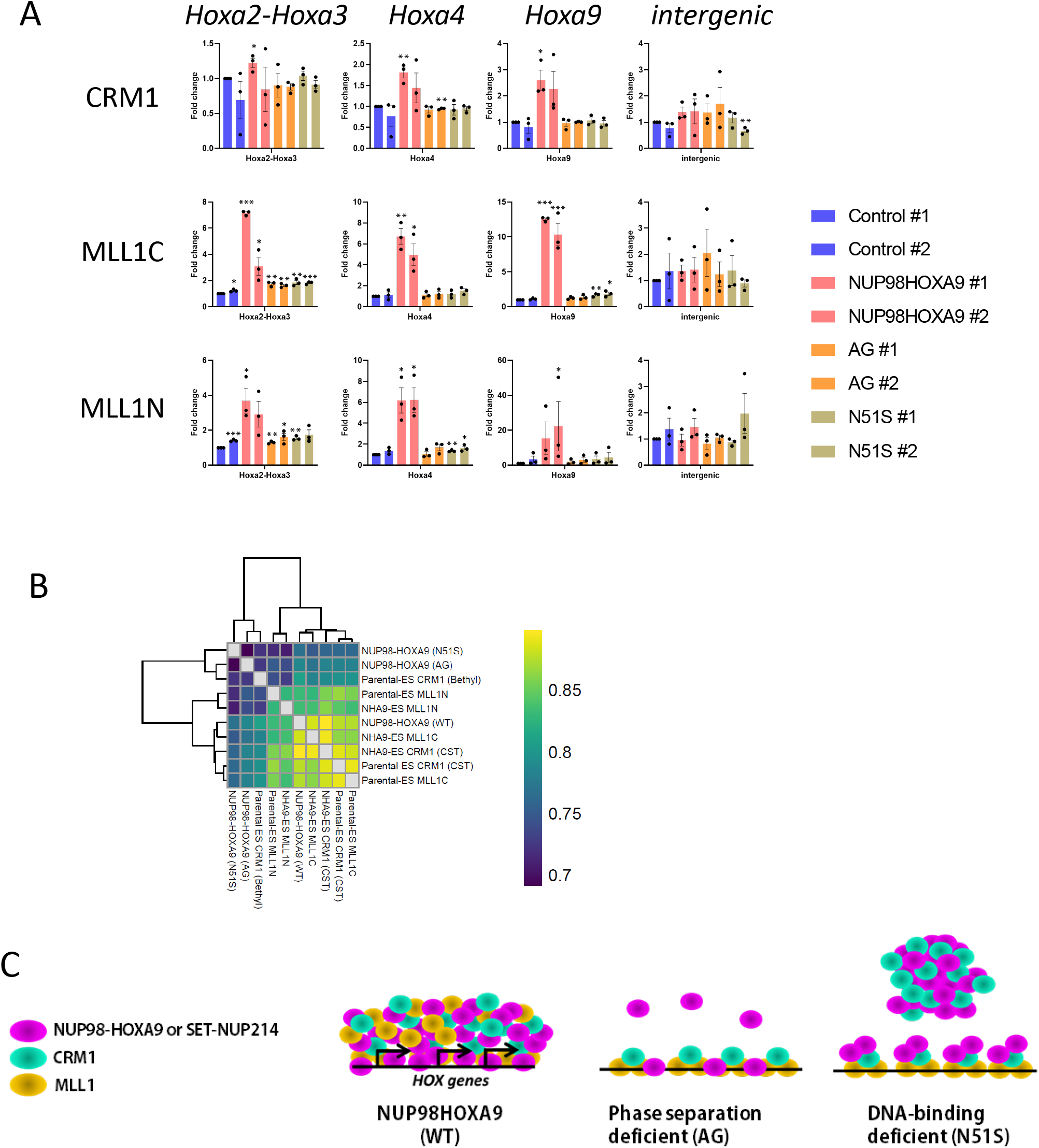
The formation of NUP98-HOXA9 nuclear bodies with an intact homeodomain is important to enhance the recruitment of CRM1 and MLL1 on *Hox* region. (A) ChIP-qPCR analysis of CRM1, MLL1N and MLL1C at Hox cluster (*Hoxa2-a3, Hoxa4, Hoxa9*) and intergenic control region (chr18) in mESCs stably expressing FLAG-NUP98-HOXA9, AG mutant, homeodomain mutant (N51S). Two independent cell lines were used. Data are presented as mean values ± SEM of three independent experiments (n = 3). Asterisks indicate statistical significance determined by Student’s *t*-test; * *p* < 0.05; ** *p* < 0.01; *** *p* < 0.001. (B) Correlation plots of MLL1N, MLL1C, and CRM1(CST) binding sites in parental mESCs (Parental-ES) or NUP98-HOXA9-expressing mESCs (NHA9-ES), together with FLAG-NUP98HOXA9, AG mutant, homeodomain mutant (N51S). (C) NUP98-HOXA9 or SET-NUP214 forms molecular condensates with CRM1 and MLL1 on its target sites including *Hox* cluster regions. In phase-separation-deficient mutant, neither the accumulation of NUP-fusion, CRM1, MLL1 are observed. In DNA-binding deficient mutant, NUP-fusion can form nuclear bodies, but those are not substantially associating with the chromatin of target genes. Note that MLL1 is not recruited in these nuclear bodies.

Together, these results showed that NUP fusion is capable of forming molecular condensates with CRM1 and MLL1, which induce and/or maintain the activation of a wide range of its targets.

## DISCUSSION

In this study, we demonstrated that two NUP fusions, SET-NUP214 and NUP98-HOXA9, drive the formation of nuclear bodies containing CRM1 and MLL1. Especially, our results showed that phase separation of NUP fusions induces the condensation of MLL1 on specific target sites.

MLL1 has been implicated in NUP fusion activity. NUP98-HOXA9 has been shown to be associated with MLL1 and is required for its recruitment to target sites, including *HOX* clusters, and downstream gene activation (Shima et al., 2017; Xu et al., 2016). Moreover, MLL1 knockout prevents NUP98-HOXA9-driven leukemogenesis (Shima et al., 2017; Xu et al., 2016). In addition, SET-NUP214 has also been demonstrated to be associated with MLL1 and to cooperatively enhance the transcription of *HOXA10* (Cigdem et al., 2021).

Here, our results suggested that NUP fusions and MLL1 not only interact with each other but they are both essential constituents of molecular condensate (i.e., nuclear bodies) that are presumably suitable for the simultaneous activation of several *HOX* genes that span tens of kb apart, as observed in leukemia cells. MLL1, a histone methyltransferase, and its associated molecules are most likely primary molecules for the creation of this molecular environment for gene activation. However, we cannot exclude the possibility that other unknown factors could also be recruited to achieve robust gene activation through phase separation. Our study further demonstrated that these NUP bodies are maintained in a CRM1-dependent manner.

How these NUP bodies function and why their NUP properties are important? In addition, the reason why only NUP98 and NUP214, among 30 different nucleoporins, are frequently found in leukemogenic fusion remains unknown. A common characteristic of NUP98 FG and NUP214 FG is the existence of dense FG repeat sequences. These FGs can form FG hydrogels *in vitro* (Labokha et al., 2013). In particular, Nup98 FG domains, which contain GLFG motifs, are highly cohesive. NUP98, even from various species, can be shown to phase separate to create FG particles (Schmidt and Gorlich, 2015). All these NUP bodies show unique features, namely they repel inert macromolecules, yet allow the entrance of NTRs associated with their cargoes.

Because CRM1 can penetrate the FG hydrogel with its cargo (Labokha et al., 2013), we speculate that NUP98FG or NUP214FG could efficiently hold CRM1 and its associated molecules (including MLL1) into the same condensates through phase separation. However, the way that CRM1 associates with MLL1 and whether MLL1 contains a functional NES remains unclear.

MLL encodes a large protein that is proteolytically cleaved into two fragments, MLL-N and MLL-C (Hsieh et al., 2003; Yokoyama et al., 2002). Interestingly, MLL1 is known to form nuclear speckles (Yano et al., 1997) in which both MLL-N and MLL-C fragments colocalize (Hsieh et al., 2003). In addition, the prediction of disordered residues using IUPred2A (Meszaros et al., 2018) suggested the presence of significant intrinsically disordered regions (IDRs) in full-length MLL1(Figure 7—figure supplement 1). This suggests that MLL1 is prone to phase separation, and its interaction with NUP fusions or CRM1 may further stimulate the formation of nuclear bodies on chromatin. In fact, the binding profiles revealed by ChIP-seq showed that NUP fusion, together with MLL1 and CRM1, binds to an unusually wide span of target gene loci. Thus, the formation of NUP-fusion nuclear bodies on chromatin seems suitable for creating a molecular environment for the simultaneous activation of multiple target genes located in the neighborhood, as represented by the *HOX-A* cluster genes.

In a recent study, Ahn et al. showed that FG-mediated LLPS (Liquid-Liquid Phase Separation) of NUP98-HOXA9 is important for the induction of an aberrant three-dimensional chromatin structure that promotes malignant transformation (Ahn et al., 2021). They observed that DNA loops were specifically formed in NUP98-HOXA9 overexpressed 293FT cells. Furthermore, they found that the loops frequently overlapped with H3K27Ac marks. However, an interactome study using BioID suggested that both wild-type and phase-separation-deficient NUP98-HOXA9 (FG to SG mutant) shared most of their interacting proteins, including MLL1. In contrast, our results demonstrated that NUP98-HOXA9, but not its phase-separation-deficient mutant nor DNA-binding deficient mutant, could induce the condensation of MLL1 around its target sites, including the *HOX-A* cluster region (Figure 7C). Collectively, the nuclear bodies formed by NUP98-HOXA9 may be important for both three-dimensional chromatin architecture as well as recruitment and/or concentration of MLL1. Future studies are needed to elucidate the relationship between molecular condensation, higher-order chromatin stricture, and gene activation, which are all triggered by the nuclear bodies of NUP fusions.

## MATERIALS AND METHODS

### Cell culture

The leukemia cell line LOUCY was cultured in RPMI1640 medium (Sigma) supplemented with 20% fetal bovine serum (FBS) (Sigma). EB3 ES cells (Niwa et al., 2002; Ogawa et al., 2004), NUP98-HOXA9 stable ES cell lines or their derivatives were cultured as described previously (Oka et al., 2016).

### RIME

RIME was performed as previously described (Mohammed et al., 2016). Briefly, 1.5 × 10^7^ LOUCY cells per condition were collected and fixed in serum-free RPMI medium supplemented with 1% (v/v) formaldehyde for 8 min at room temperature. After the addition of a final 0.1 M glycine to quench crosslinking, cells were washed twice with ice-cold phosphate-buffered saline (PBS) and resuspended in 500 μL of PBS. After removal of the supernatants, the cell pellets were snap-frozen in liquid N2 and stored at −80 °C. The nuclear fraction of the cells was extracted by resuspending the pellet in 10 mL of LB1 buffer (50 mM HEPES-KOH [pH 7.5], 140 mM NaCl, 1 mM EDTA, 10% glycerol, 0.5%Igepal CA-630, and 0.25% Triton X-100) for 10 min at 4 °C with rotation. The cells were collected and resuspended in 10 mL of LB2 buffer (10 mM Tris-HCl [pH 8.0], 200 mM NaCl, 1 mM EDTA, and 0.5 mM EGTA) and incubated at 4 °C for 5 min with rotation. Cells were pelleted and resuspended in 300 mL of LB3 buffer (10 mM Tris-HCl [pH 8], 100 mM NaCl, 1 mM EDTA, 0.5 mM EGTA, 0.1% Na-deoxycholate, and 0.5% N-lauroylsarcosine) and sonicated in a water bath sonicator (Diagenode Bioruptor; on ice, 30 s on/off cycle for 20 min). After the addition of 30 μL 10% (v/v) Triton X-100, the samples were vortexed and centrifuged at 20,000 x g for 10 min to remove cell debris. The supernatant was then added to 100 μL of magnetic beads (Dynal), preincubated with antibody, and immunoprecipitation (IP) was performed at 4 °C overnight. The beads were then washed with 1 mL of RIPA buffer 10 times and sequentially washed with freshly prepared 100 mM ammonium hydrogen carbonate (AMBIC) solution. For the second AMBIC wash, beads were transferred to new tubes. After removing the supernatant, the samples were stored at – 80 °C.

To prepare the magnetic beads bound to the antibody, 100 μL of magnetic beads (Dynal) was resuspended in 1 mL of PBS w/5 mg/mL BSA (PBS/BSA) and washed four times with a magnetic separator. After resuspension in 500 μL of PBS/BSA, each antibody was added and incubated overnight with rotation. The next day, the beads were washed five times with PBS/BSA and used for IP.

Bead-bound protein was digested with 100 ng trypsin (Promega, V5113) overnight at 37 °C. After the overnight digest, 100 ng of trypsin was added to each sample and digested for 4 h at 37 °C. The digested peptides were acidified with TFA, desalted, and purified on C18-SCX StageTips (Adachi et al., 2016). The peptides were dried with SpeedVac and solubilized with 0.1% formic acid/2% acetonitrile.

### Liquid chromatography with tandem mass spectrometry (LC-MS/MS) analysis

LC-MS/MS was performed by coupling an UltiMate 3000 Nano LC system (Thermo Fisher Scientific) and HTC-PAL autosampler (CTC Analytics) to a Q Exactive hybrid quadrupole-Orbitrap mass spectrometer (Thermo Fisher Scientific). The peptides were delivered to an analytical column (75 μm × 30 cm, packed in-house with ReproSil-Pur C18-AQ, 1.9 μm resin, Dr. Maisch, Ammerbuch, Germany) and separated at a flow rate of 280 nL/min using a 145 min gradient from 5% to 30% of solvent B (solvent A, 0.1% FA and 2% acetonitrile; solvent B, 0.1% FA, and 90% acetonitrile). The Q Exactive instrument was operated in data-dependent mode. Survey full-scan MS spectra (*m/z* 350–1,800) were acquired in the Orbitrap with 70,000 resolution after the accumulation of ions to a 3 × 10^6^ target value. Dynamic exclusion was set at 30 s. The 12 most intense multiplied charged ions (z ≥ 2) were sequentially accumulated to a 1 × 10^5^ target value and fragmented in the collision cell by higher energy collisional dissociation (HCD) with a maximum injection time of 120 ms and 35,000 resolution. Typical mass spectrometric conditions were as follows: spray voltage, 2 kV; heated capillary temperature, 250 °C; normalized HCD collision energy, 25%. The MS/MS ion selection threshold was set to 2.5 × 10^4^ counts. A 2.0 Da isolation width was chosen.

### Data processing and visualization

Raw MS data were processed using MaxQuant (version 1.6.14.0), supported by the Andromeda search engine. The MS/MS spectra were searched against the UniProt human database with the following search parameters: full tryptic specificity; up to two missed cleavage sites; carbamidomethylation of cysteine residues set as a fixed modification; serine, threonine, and tyrosine phosphorylation, N-terminal protein acetylation; and methionine oxidation as variable modifications. The false discovery rate of the protein groups and peptides was less than 0.01. Peptides identified from the reversed database or those identified as potential contaminants were not used.

### ImmunoFISH

ImmunoFISH was performed as previously described (Chaumeil et al., 2013). LOUCY cells were pelleted, resuspended in a small volume of PBS, and placed on a poly L-lysine-coated coverslip (Neuvitro Corporation, H-15-PDL). After the fixation of cells in 2% paraformaldehyde/1x PBS for 10 min at room temperature, cells were washed three times in PBS and permeabilized with ice-cold 0.4% Triton-X-100 in PBS for 5 min on ice. Then, the cells were incubated in 100 μL of blocking solution for 30 min at room temperature and incubated with the primary antibody in 100 μL of blocking solution for 1 h at room temperature. After washing the cells in 0.2% BSA/0.1% Tween-20/PBS three times for 5 min each, the cells were incubated with the secondary antibody diluted in 100 μL of blocking solution for 1 h. The cells were washed three times with 0.1% Tween-20/PBS for 5 min each and fixed in 2% paraformaldehyde/PBS for 10 min. After three washes with PBS, cells were incubated with 0.1 μg/μL RNase A (Nippon Gene) in PBS for 1 h at 37 °C in FISH slide processing system (Thermobrite, ABBOTT). Cells were washed with PBS three times and permeabilized in ice-cold 0.7% Triton-X-100/0.1 M HCl for 10 min on ice. After three washes with PBS, the cells were incubated in 50% formamide/2x SSC at 75 °C for 30 min and hybridized with pre-labelled BAC probes (RPCI-11 29G13 labelled red for *HOX-B*(chr17) and RPCI-11 44L12 labelled green for *RNF2*(chr1); Empire Genomics) overnight at 37 °C using FISH slide processing system. The next day, the cells were washed three times in 50% formamide/2x SSC, 5 min each at 37 °C, followed by another three washes in 2x SSC (the last wash was performed with 2x SSC containing 4’,6-diamidino-2-phenylindole (DAPI), 5 min each at 37 °C. Cells were mounted in ProLong Gold mounting medium (Thermo Fisher Scientific), and observed using a SP8 confocal microscope (Leica). To evaluate the distance between SET-NUP214 and each genomic locus, z-stack confocal images of SET-NUP214 and FISH were projected in 2D using max z projection function in imageJ. Projecected images were binarized and NUP214 particles were registed by imageJ roi manager. The distance from gene loci were then calculated using distance map function of imageJ. The closest distance from gene loci were then calculated by measuring the minimum intensity in the distance map.

### Plasmids, isolation of stable ES cell lines

Stable ES cell lines were generated as previously described (Oka et al., 2016). Briefly, parental ES cells were transfected with pCAGGS-m2TP, an expression vector for Tol2 transposase, and pT2A-CMH-FLAG-NUP98-HOXA9 or its mutants (Kawakami and Noda, 2004; Urasaki et al., 2006) using Lipofectamine 2000 (Life Technologies). After 2 days, the cells were trypsinized and replated onto ES-LIF medium containing hygromycin B (200 μg/mL). Colonies were picked, and the expression of FLAG-NUP98-HOXA9 or its mutants was examined by immunoblotting and immunofluorescence staining. cDNA encoding NUP98-HOXA9 has been previously described (Oka et al., 2016), whereas DNA encoding the AG mutant of NUP98-HOXA9 was synthesized (GenScript)(supplememntal file 1), or the homeodomain (N51S) mutant of NUP98-HOXA9 was generated by PCR-based mutagenesis, and cloned into pT2A-CMH-FLAGx3.

### Immunostaining and confocal microscopy

Human leukemia LOUCY cells were attached onto poly L-lysine-coated coverslips (Neuvitro Corporation, H-15-PDL) and fixed in PBS containing 3.7% formaldehyde for 15 min at room temperature. The cells were washed with PBS and permeabilized with 0.5% Triton X-100 in PBS for 5 min at room temperature. After washing with PBS, the cells were incubated with blocking buffer (3% skim milk in PBS) for 30 min. The cells were incubated with primary antibodies overnight, washed three times with PBS, and incubated with secondary antibodies for 30 min. The cells were washed twice with PBS and stained with DAPI in PBS for 15 min at room temperature. The cells were mounted using ProLong Gold (Thermo Fisher Scientific). Images were acquired using a SP8 confocal microscope (Leica). ES cells were grown on coverslips and fixed in a medium containing 3.7% formaldehyde for 15 min at room temperature. After washing with PBS, the cells were permeabilized with 0.5% Triton X-100 in PBS for 5 min and further incubated with a blocking buffer (3% skim milk in PBS) for 30 min. Next, cells were incubated with primary antibodies overnight at 4 °C. After washing four times with PBS, the cells were incubated with secondary antibodies for 30 min. Cells were washed four times with PBS, stained with DAPI for 15 min at room temperature, and the coverslips were mounted with ProLong Gold mounting medium (Thermo Fischer Scientific). Images were acquired using a SP8 confocal microscope (Leica).

### ChIP-qPCR, ChIP-seq

ChIP-qPCR and ChIP-seq were performed as previously described (Oka et al., 2019). Human leukemia LOUCY cells were fixed in a medium containing 0.5% formaldehyde at room temperature for 5 min. Fixed cells were collected by centrifugation at room temperature, washed twice with ice-cold PBS, and resuspended in ChIP buffer (10 mM Tris-HCl pH 8.0, 200 mM KC), 1 mM CaCl2, 0.5% NP40) containing protease inhibitors (2 μg/mL aprotinin, 2 μg/mL leupeptin, and 1 μg/mL pepstatin A). Cells were briefly sonicated (Branson 250D Sonifier, Branson Ultrasonics), and after centrifugation, the supernatants were digested with 1 U/mL micrococcal nuclease (Worthington Biochemical) for 40 min at 37 °C. The reaction was stopped with EDTA (final concentration of 10 mM), and the supernatants were incubated with anti-mouse or anti-rabbit IgG magnetic beads (Dynabeads, Life Technologies) preincubated with anti-FLAG M2 (Sigma), anti-CRM1 (CST or Bethyl Lab), anti-NUP214 (Bethyl Lab), anti-MLL1C (CST), and anti-MLL1N (CST) for 6 h. The beads were washed twice with each of the following buffers: ChIP buffer, ChIP wash buffer (10 mM Tris-HCl pH 8.0, 500 mM KCl, 1 mM CaCl2, 0.5% NP40), and TE buffer (10 mM Tris-HCl pH 8.0, 1 mM EDTA) and eluted in an elution buffer containing 50 mM Tris-HCl pH 8.0, 10 mM EDTA, and 1% sodium dodecyl sulfate overnight at 65 °C. DNA was recovered using AMPure XP beads (Beckman Coulter) and used for ChIP-qPCR analysis or for library preparation for ChIP-seq analysis.

For ES cells, cells grown in dishes were fixed in medium containing 0.5% formaldehyde at room temperature for 5 min. After one wash with ice-cold PBS, PBS (0.5 mL) was added, and cells were collected with a scraper. After centrifugation, the collected fixed cells were processed as described above for LOUCY cells.

The ChIP library was prepared using ThruPLEX DNA-seq kit (Takara Bio), according to the manufacturer’s instructions, and sequenced on the Illumina HiSeq1500 or NovaSeq 6000 system. Adaptor sequence and low quality sequence were removed and read length below 20bp were discarded by using Trim Galore (version 0.6.70). The sequence reads were aligned to the reference mouse genome (mm10) and human genome (GRCh38) using Bowtie2 (version 2.3.1). Multi-mapping and duplicated reads were not used for further analysis. ChIP peaks were indentified by using MACS2 (version 2.2.7.1) with p-value < 0.001.

### qPCR

Total RNA was extracted from cells using the MagMAX mirVana Total RNA kit (Thermo Fisher Scientific) using King Fischer Duo (Thermo Fisher Scientific) or ReliaPrep RNA Miniprep Systems (Promega) and used for cDNA synthesis with the PrimeScript RT reagent Kit (Takara Bio). All procedures were performed in accordance with the manufacturer’s protocol. qPCR analysis was performed on a 384-well plate with the QuantStudio 6 Flex Real-Time PCR System (Life Technologies) using GeneAce SYBR qPCR Mix (Nippon Gene). The relative gene expression levels were normalized to *GAPDH* mRNA levels as a control. The primer sequences are listed in Supplementary File 1.

### Gene knockdown

Gen knockdown in LOUCY cells was performed as previously described (Oka et al., 2019). Briefly, LOUCY cells (1 × 10^7^ cells) were transfected with 4 μM of each siRNA (TriFECTa® RNAi Kit, IDT) or negative control siRNA (Universal Negative Control siRNA [Nippon Gene]) by nucleofection (Lonza) using reagent V and the X-001 program. Immediately after nucleofection, the cells were plated on a 100 mm dish in RPMI 1640 medium supplemented with 20% FBS for the indicated period and used for immunoblotting and ChIP-qPCR.

### Accession numbers

The ChIP-Seq data are accessible through GEO Series accession number GSE202837.

### Statistical analysis

Statistical analyses were performed using an unpaired two-tailed Student’s *t*-test. GraphPad Prism version 8 was used for data analysis and representation.

## ACKNOWLEDGEMENTS

We thank Dr. Koichi Kawakami for transposon vectors, Dr. Hitoshi Niwa for ES cells. This work was partly performed in the Cooperative Research Project Program of the Medical Institute of Bioregulation, Kyushu University.

---

**Figure 1—figure supplement 1.**
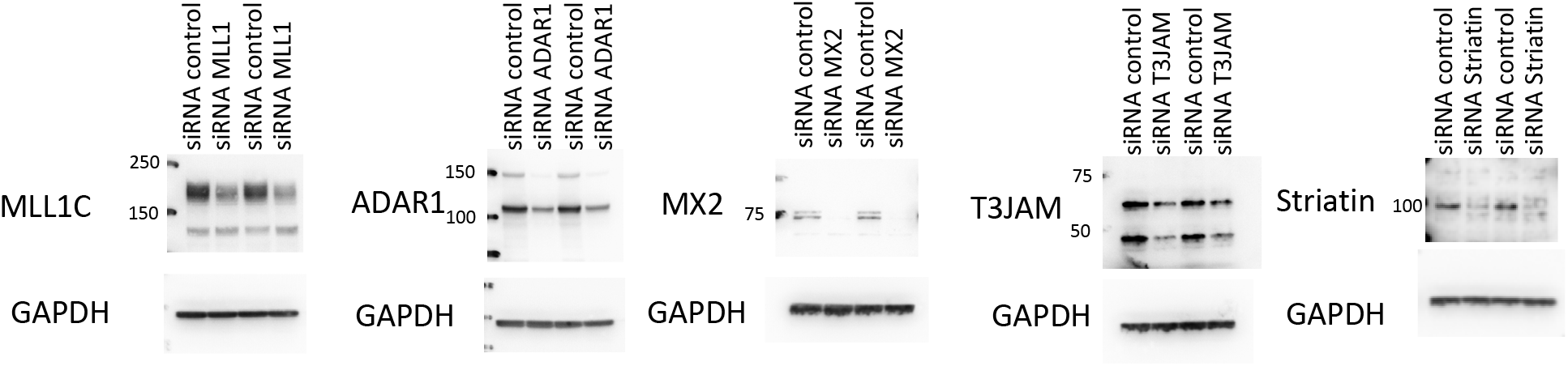
Validation of knockdown by western blot analysis. Knockdown of indicated proteins were examined 4 days after nucleofection with either negative control siRNA or siRNA (DsiRNA, IDT) against targets by immunoblotting using indicated antibodies.

**Figure 6—figure supplement 1.**
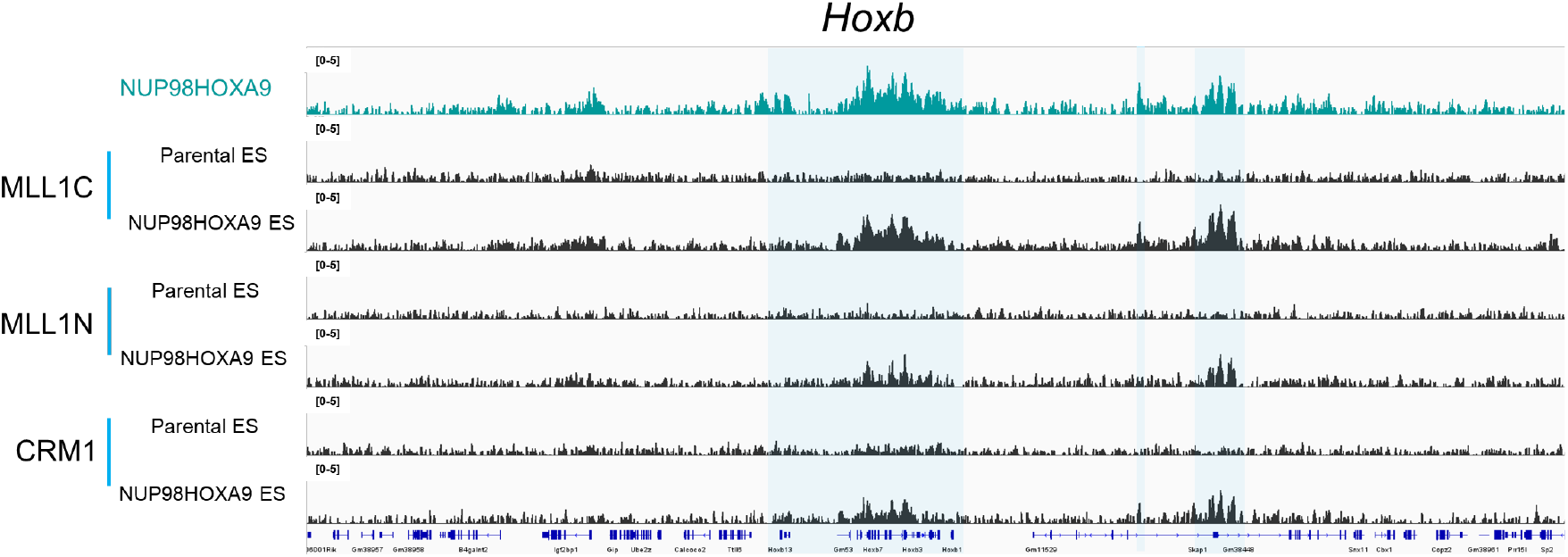
NUP98-HOXA9 nuclear bodies stimulate the condensation of MLL1 on *Hoxb* cluster. The binding profiles of MLL1C, MLL1N, CRM1 on *Hoxb* cluster in parental mESC or stable cell lines expressing FLAG-NUP98-HOXA9 as revealed with ChIP-seq.

**Figure 7—figure supplement 7.**
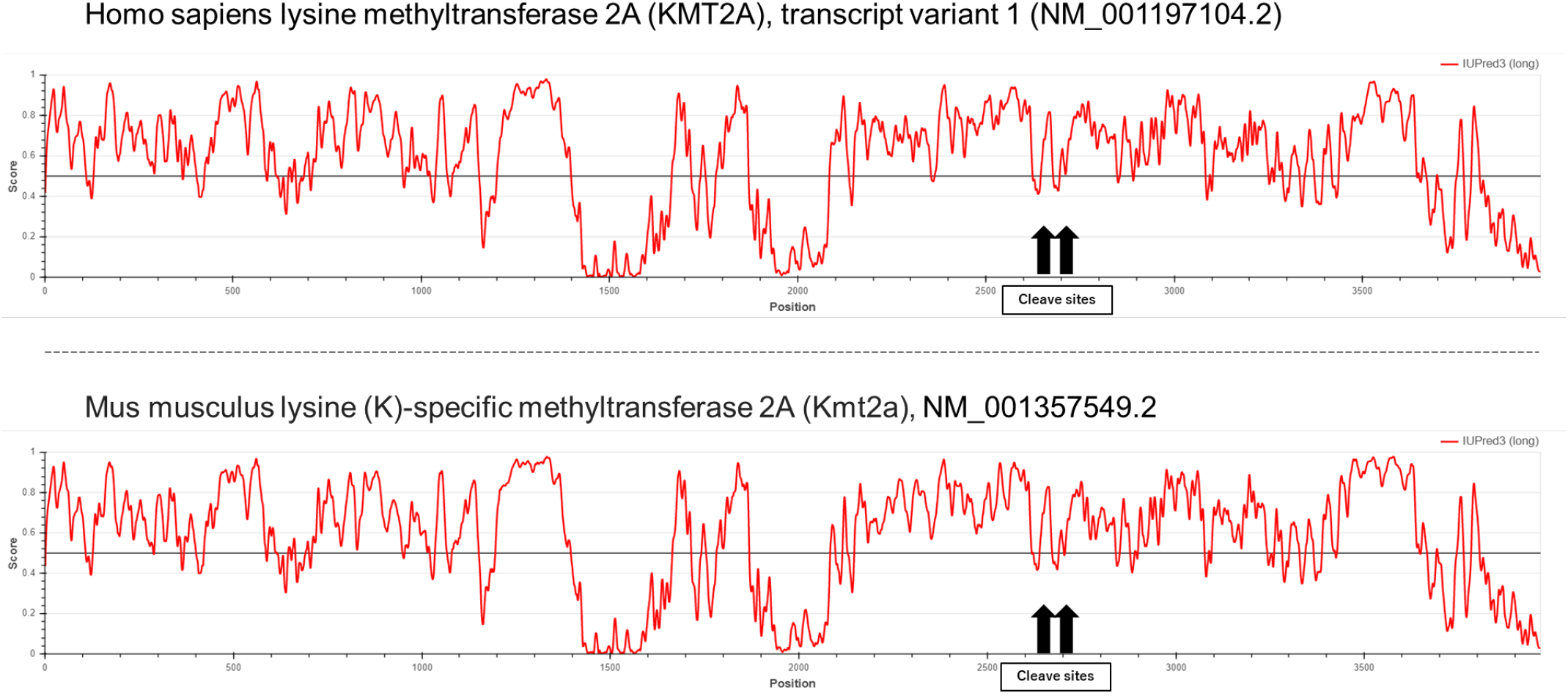
The intrinsically disordered regions of MLL1s are predicted using IUPred3A (intrinsically unstructured/disordered proteins and domains prediction tool). The cleavage sites of MLL1 are shown by arrows.

